# Comparing the outputs of intramural and extramural grants funded by National Institutes of Health

**DOI:** 10.1101/2023.11.09.566298

**Authors:** Xiang Zheng, Qiyao Yang, Jai Potnuri, Chaoqun Ni, B. Ian Hutchins

## Abstract

Funding agencies use a variety of mechanisms to fund research. The National Institutes of Health in the United States, for example, employs scientists to perform research at its own laboratories (intramural research), and it also awards grants to pay for research at external institutions such as universities (extramural research). Here, using data from 1594 intramural grants and 97054 extramural grants funded between 2009 and 2019, we compare the scholarly outputs from these two funding mechanisms in terms of number of publications, relative citation ratio and clinical metrics. We find that extramural awards are more cost-effective for producing outputs commonly used for academic evaluation, such as publications and citations (per dollar), while intramural awards are more cost-effective for generating research that influences future clinical work, more closely in line with the agency’s health goals. These findings provide evidence that institutional incentives associated with different funding mechanisms drive their comparative strengths.

## Introduction

National funding agencies have a responsibility to ensure that the research projects they fund are successful, as measured by various metrics (such as number of scientific papers, number of early-career researchers trained, and societal and/or economic impact). Many of these agencies run their own research laboratories (intramural research) and also fund research at universities and other institutions (extramural research). Certain aspects of the grant funding system have been the focus of research, such as publication of highly influential papers (Azoulay, Zivin, & Manso, 2009) risk management (Goldstein & Kearney, 2020), funding disparities (T. A. Hoppe et al., 2019) and diminishing marginal returns to funding directed to single labs (Lauer, Roychowdhury, Patel, Walsh, & Pearson, 2017; Wahls, 2018a, 2018b) However, the relative merits of intramural and extramural funding have received little attention to date.

The United States National Institutes of Health is one of the largest funders of research in the world. It comprises 27 institutes and centers (24 of which fund grants), each with its own research agenda often related to a specific disease (such as the National Cancer Institute and the National Institute of Allergy and Infectious Diseases). In Fiscal Year 2022, the NIH spent approximately $5 billion on intramural research at its own laboratories, and $39 billion on extramural research at universities, medical schools and research institutes across the US (“NIH Budget,” 2025). Most applications for extramural grants are peer reviewed and assigned a percentile ranking of overall impact merit score by a Study Section (“What happens to your application during and after review?,” 2025), with the relevant NIH institute or center making a final decision on which applications are funded based on those scores. Intramural research is conducted in government laboratories run by Senior Investigators (Sampat, 2012), supported by a combination of staff scientists and postdoctoral fellows. Senior investigators do not have to apply for grants, but external Boards of Scientific Counselors review their performance on a regular basis (usually every four years). Questions about the most effective portfolio management approaches have been an ongoing source of contention (Supplemental Text).

A potential advantage of the extramural approach is that cost-sharing at universities may increase the scientific return on the NIH’s investment. An advantage of the intramural approach are that NIH has the direct ability to hire scientists whose research closely aligns with agency goals, and researchers do not need to devote time and effort on preparing and submitting grant applications. Here, using data from 1594 intramural grants and 97054 extramural grants, we compare the distribution of research topics funded by the two mechanisms (Figure 1), and the values of the grants awarded under each mechanism (Figure 2). The large difference in the number of intramural vs. extramural awards is reflective of the larger size of intramural awards compared to extramural awards, combined with the much smaller proportion of the funding portfolio dedicated to intramural research. We also use five metrics to analyze the 621,138 papers that acknowledge at least one of these projects: number of papers; relative citation ratio, which is a field- and time-normalized measure of scientific influence; approximate potential to translate, which is a machine-learning prediction that a given paper will be cited by a clinical article; total clinical citation counts; and a binary measure of the number of papers that received at least one clinical citation (Figure 3). We also compare the cost effectiveness of the two approaches by, for example, comparing the average cost of each paper published (Figure 4), and also explore the influence of various factors, such as the high costs associated with human-focused research (Figure 5).

**Figure 1.**
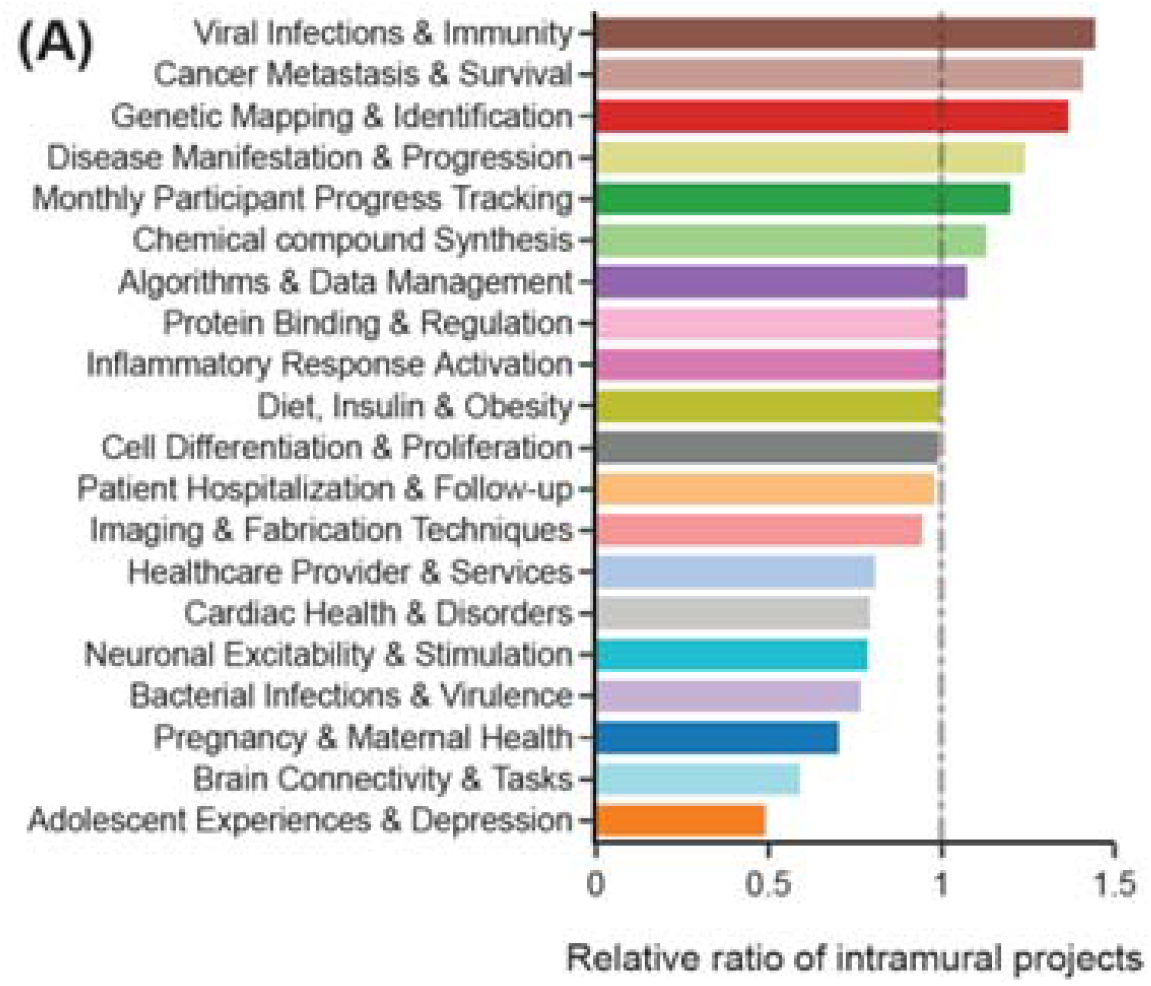
Research topics for intramural and extramural projects. The topics listed were identified by clustering projects based on their titles and abstracts via Word2Vec (see Methods). The relative ratio of intramural projects for each topic was calculated by taking a ratio of the proportions of total grants a topic represented in the intramural vs. extramural portfolios. A relative ratio >1 signifies a higher share of intramural projects on that topic relative to their share across all topics. For example, if a topic comprised 10% of grants in the intramural portfolio but only 5% of grants in the extramural portfolio, this would represent a 2:1 intramural:extramural relative ratio, or 2.0.

**Figure 2.**
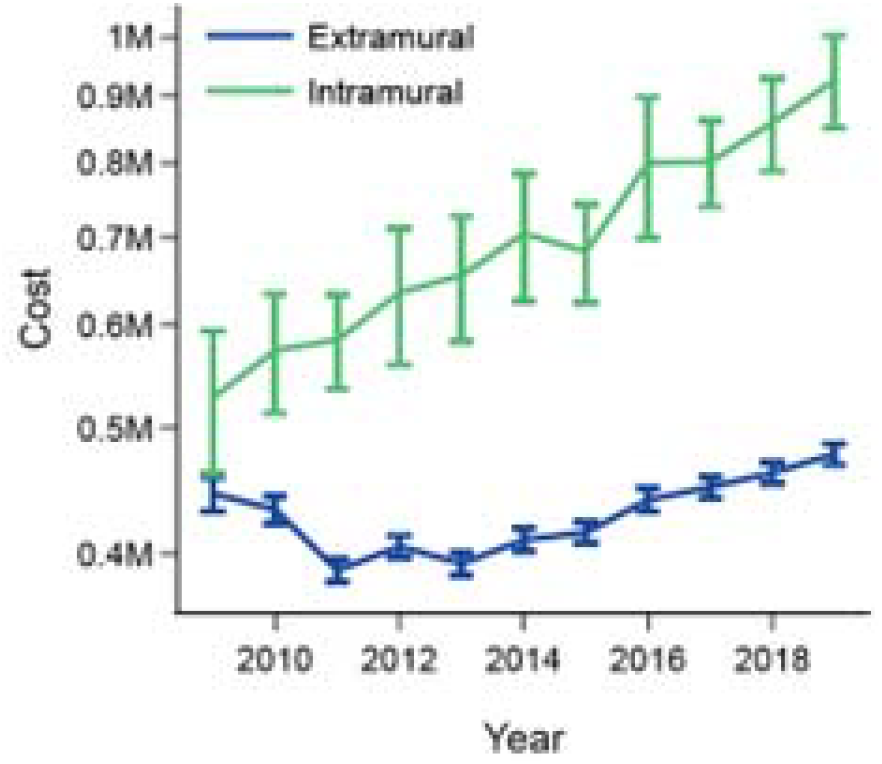
Project funding for intramural and extramural projects. Mean project cost (on a log scale) versus year for intramural grants (green) and extramural grants (blue) between 2009 and 2019. Error bars denote 95% confidence intervals. Total costs were used rather than only direct costs in order to fully account for the degree of government investment. Error bars are larger for intramural data because of the smaller total number of awards (98,648 extramural and 1594 intramural).

**Figure 3.**
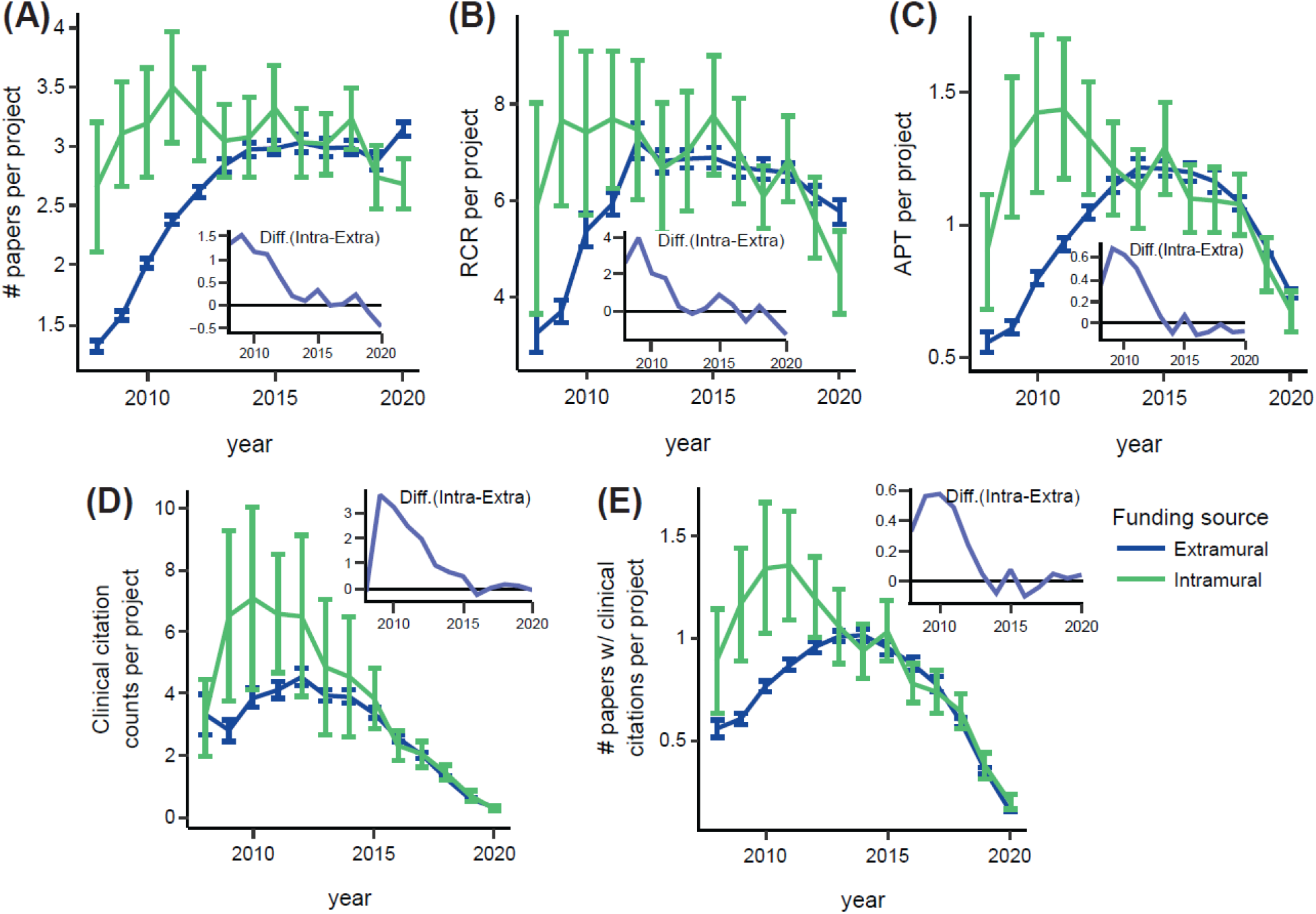
Annual outputs from intramural and extramural projects. **(A)** Mean number of papers per project for intramural projects (green) and extramural projects (blue) between 2008 and 2020. The difference (inset) was close to 1.5 papers per project in 2008, but this gap closed over time. **(B)** Relative citation ratio per project. **(C)** Approximate potential to translate (ATP) per project. **(D)** Clinical citation counts per project. **(E)** Number of papers with at least one clinical citation per project. Error bars denote 95% confidence intervals. Error bars are larger for intramural data because of the smaller total number of awards (98,648 extramural and 1594 intramural).

**Figure 4.**
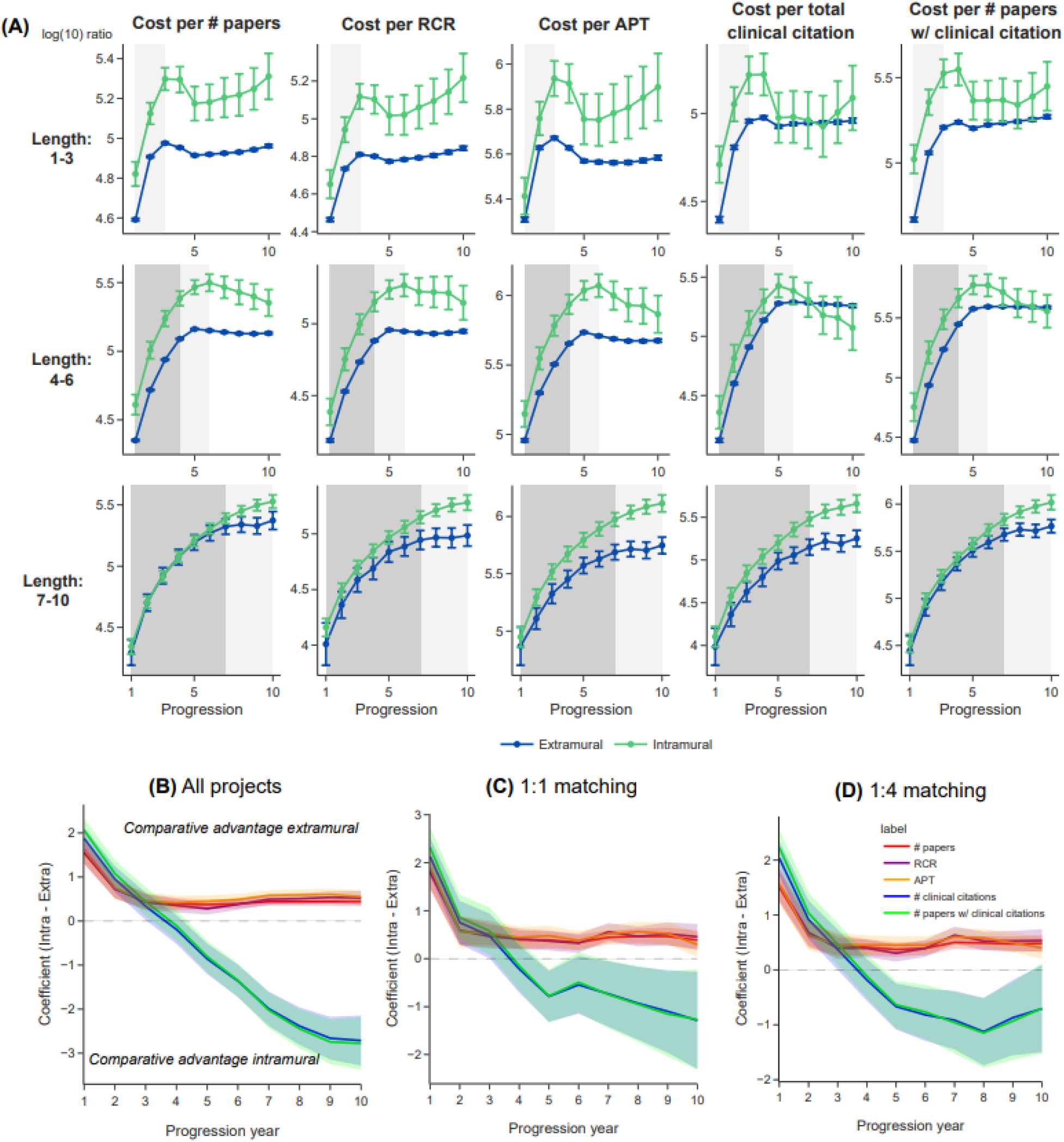
Cost effectiveness of intramural and extramural projects. **(A)** A measure of cost effectiveness versus progression (i.e., year of grant) for intramural research (green) and extramural research (blue), for projects of different durations: 1–3 years (top row), 4–6 years (middle), and 7–10 years (bottom). These regressions do not control for other characteristics, but rather represent the raw ratios. For the first column, the Y-axis displays log_10_(ratio) +1, where ratio is the cumulative total costs to the cumulative total research output for each metric (cost:output, for the first column output = #papers); error bars denote the 95% confidence intervals. The remaining columns show measures of cost effectiveness for relative citation ratio, approximate potential to translate, total clinical citation counts, and a binary measure of clinical citations. To account for the fact that many papers are published after funding for the relevant grant has ended, grant amounts were multiplied by a deflator – this represents the proportion of papers published to date against the anticipated number of future publications, as determined by empirical measurements (Supplemental Table 1). In most cases, according to this analysis, extramural research is more cost effective than intramural research when observing uncontrolled regressions. **(B–D)** Linear regression results of the cost efficiency of research output measures against project types (intramural vs. extramural). The regression model was fitted for each year of the project’s progression. Unlike panel (A), this regression model controls for grant, investigator, and collaboration characteristics in order to obtain a more accurate estimate of the relative cost efficiency of intramural vs. extramural projects. The Y-axis coefficient indicates the mean disparity in research output between intramural and extramural projects, controlling for these other variables (see Methods). Because there might be covariates that could confound the data, separate regressions were conducted for all projects (B, the default), and for balanced projects using 1:1 propensity score matching (1 extramural grant for every 1 intramural grant) in order to compare grants that were the most similar to reduce the influence of unobserved covariates (C) and (D) similarly to (C) 1:4 propensity matching as a robustness check.

**Figure 5.**
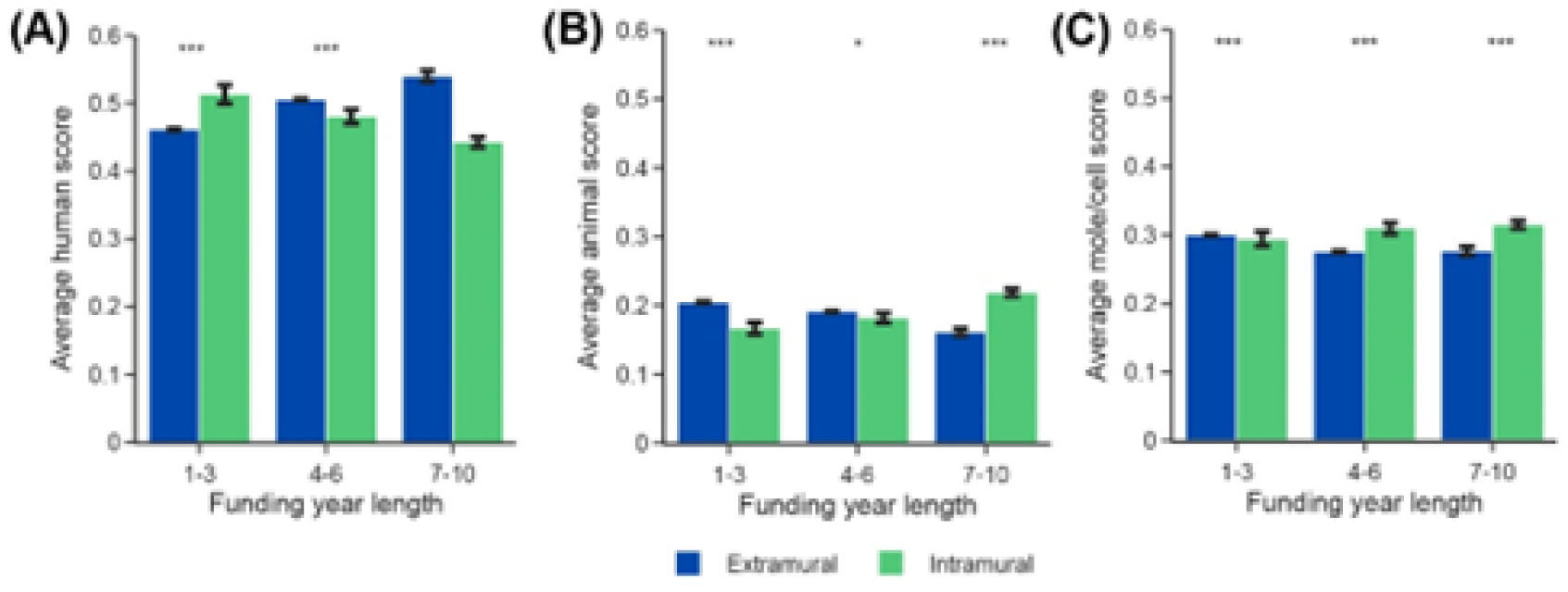
Comparison of scores for human-focused research, animal research, and molecular/cellular research for intramural and extramural projects. **(A-C)** These represent the average Human, Animal and Molecular/Cellular scores for publications funded by extramural vs. intramural grants, respectively, which were downloaded from iCite (B. I. Hutchins, Davis, et al., 2019; iCite et al., 2019). **(A)** Average scores for human-focused research for intramural research (green) and extramural research (blue) for projects of different durations: 1–3 years (left), 4–6 years (middle), and 7–10 years (right).. **(B)** Average scores for animal research. **(C)** Average scores for molecular/cellular research. Mann-Whiteney U tests were conducted to test the difference between the scores for intramural and extramural projects. *** p<0.001, ** p < 0.01, * p < 0.05. No asterisk indicates the difference is not statistically significant.

## Results

### Comparison of research topics for intramural and extramural projects

Our analyses reveal differences in the research topics investigated by researchers funded by the two mechanisms. By clustering projects based on their titles and abstracts, we find that intramural projects yield a higher-than-average number of projects on viral infection, cancer and genes, and a lower-than-average number of projects on adolescents, brain studies and maternal health (Figure 1 and Supplemental Figures 1 and 2). The overrepresentation of viral research is likely because of the outsize investment toward the intramural Vaccine Research Center, and the cancer/genetics overrepresentation due in part because National Cancer Institute intramural investigators conduct research at that institute as well as at the NIH Clinical Center and Center for Cancer Research for their human genetics work (“National Cancer Institute (NCI) Center for Cancer Research,” 2025). Intramural projects begin in our dataset with a lower proportion of the total projects considered (Supplemental Figure 3). This is because projects that were matched in 2008 were excluded as being possible continuations of existing intramural projects. Because these new projects are more typically associated with hiring of new principal investigators than in the extramural program, where established investigators can apply for a new grant at any time, the ramp-up of these intramural projects occurred more slowly.

### Comparison of funding for intramural and extramural projects

Next we compared funding for intramural and extramural research projects, and found that annual funding for extramural research projects was consistently lower than that for intramural research projects (Figure 2). For all comparisons, we use inflation-adjusted total costs, which for extramural grants includes indirect as well as direct costs. Indeed, the average funding for extramural projects remained roughly constant, at below $500k per year between 2009 and 2019, whereas the average funding for intramural projects increased from about $0.42 million to about $0.45 million over the same period. Given that NIH funding for intramural research has remained relatively constant as a percent of total funding over the years, this indicates larger single awards for intramural research while extramural investigators may increasingly require multiple concurrent grants to sustain their labs.

This finding is consistent with the intramural research requiring higher financing due to the absence of cost-sharing with universities. It also reflects observations that extramural researchers need multiple federally funded grants to sustain a lab. While such institutions do receive audited indirect costs to cover overhead associated with research (Culliton, 1992), recent research indicates that significant cost-sharing greatly influences scientific productivity. Specifically, the extent of universities subsidies for student labor costs is a direct factor in university productivity (Zhang, Wapman, Larremore, & Clauset, 2022). In contrast to extramural awards, intramural funding as reviewed by the Board of Scientific Counselors and each Institute’s scientific Director, can in principle fund an entire lab through a single, larger award. This frees the intramural investigators from the time commitment of securing grants. However, running in-house labs does entail that there is no cost sharing with external institutions, potentially raising the cost of such research (Culliton, 1992; Korn, 2015; Macilwain, 1999; Zhang et al., 2022).

### Comparison of outputs from intramural and extramural projects

Next we used five metrics related to publications and citations – number of papers; relative citation ratio (Arabi, Ni, & Hutchins, 2025; B. I. Hutchins, Hoppe, Meseroll, Anderson, & Santangelo, 2017; B. I. Hutchins, Yuan, Anderson, & Santangelo, 2016); approximate potential to translate (B. I. Hutchins, Davis, Meseroll, & Santangelo, 2019); total clinical citation counts (B. I. Hutchins, Baker, et al., 2019; B. I. Hutchins, Davis, et al., 2019; iCite, Hutchins, & Santangelo, 2019); and a binary measure of clinical citations – to compare the outputs of intramural and extramural projects on a year-by-year basis (Figure 3). For all of the metrics apart from total clinical citation counts, intramural research scored highest before 2010, but the gap between intramural and extramural research had closed by 2020. This increased early productivity for intramural projects may reflect the extra time intramural investigators save because they do not have teaching and grant writing responsibilities. Because this dataset is truncated (e.g. only papers and their citations through 2020 are considered), we observe a decrease in most outcome measures in Figure 3 as the end of this window approaches. In our previous study (B. I. Hutchins, Davis, et al., 2019), we observed a lag of approximately seven years for clinical citations to accrue. We see the same trend in this dataset (Figure 3 D-E), approaching zero clinical citations as our time window approaches. The Approximate Potential to Translate metric, a prediction of future clinical citations, also decreases but to a lesser extent. This is because the forward citation network is used for making these predictions.

### Comparison of cost effectiveness for intramural and extramural projects

So far we have seen that the average funding for intramural projects is higher than that for extramural projects (Figure 2), and that intramural projects also score higher than extramural projects on the five metrics for the outputs from a project that we computed (Figure 3). Next, therefore, we compare cost effectiveness by calculating the average cost of each published paper, and likewise for the other four publication/citation metrics (Figure 4A). We do this for projects of different durations: 1–3 years, 4–6 years, and 7–10 years.

Our analysis suggests that extramural research is more cost-effective than intramural research when considering number of papers, relative citation ratio and approximate potential to translate. However, when considering metrics based on clinical citations, the gap was smaller, and for some project durations, intramural research was as cost-effective as extramural research.

Some types of research are more expensive rather others (for example, animal research is more expensive than cell biology research, and human-focused research has higher regulatory overhead costs due to increased ethical concerns), so we decided to explore if such factors could explain some of the differences that we had observed between intramural and extramural projects. To do this we conducted a regression analysis that controlled for project topic and various factors related to the principal investigator who had received the grant.

Figure 4B shows a plot of the number of years that elapsed since the start of the grant (Progression year, x-axis) and the relative cost effectiveness of intramural vs. extramural grants at funding each metric measured (#papers, red; RCR, purple; APT, yellow; #clinical citations, blue; and #papers with at least one clinical citation, green). Each curve was a subset of the data frame focusing on each measure individually. These were generated by comparing the difference in regression coefficients of extramural vs. intramural research once controlling for grant, investigator and collaboration variables (for regression model details, see Methods). Curves above 1.0 indicate a comparative cost effectiveness advantage for extramural grants on that measure, while those below 1.0 indicate a comparative cost effectiveness advantage for intramural grants on that measure. We observe that over time, extramural and intramural grants show opposite patterns. Extramural grants excel in terms of cost advantage at measures commonly used in academic performance assessment (e.g. number of papers and citations). Intramural grants in comparison excel in cost advantage in measures more closely aligned with the NIH agency mission (i.e. knowledge that informs work to improve human health via knowledge flow to clinical trials).

We also performed propensity score matching between intramural and extramural projects by pairing each intramural project with its closest extramural counterpart (Figure 4C), and with its four closest extramural counterparts (Figure 4D). This allows us to sample intramural and extramural grants that are similar to one another at a 1:1 (e.g. 1594 intramural grants and 1594 matched extramural grants) based on their similarity in terms of topic area, prior publication record, and prior collaboration history. This propensity score matching approach accounts for potentially hidden covariates and reduces their influence. We observe the same pattern using 1:1 propensity score matching (Figure 4C) as we did with the full dataset (Figure 4B). Likewise, using propensity score matching of 1:4 (i.e. 4 matched extramural grants for every 1 intramural grant) to increase the sample size of extramural awards yielded the same result (Figure 4D). Taken together, these results indicate that regardless of sampling strategy, cost effectiveness aligns with the primary missions of the institutions at which investigators are housed (maximum knowledge generation and flow for academic institutions, and clinically relevant knowledge generation for NIH intramural research).

It is estimated that universities do not fully recover expenditures through indirect costs. The magnitude of this effect is estimated to be approximately 30% (Droegemeier, 2017). We therefore asked whether these trends hold when indirect costs are inflated by the same amount, to more completely reflect research investment expenditures. Supplemental Fig 4 shows that inflating indirect costs by 30% narrows the gap in estimated costs between intramural and extramural awards as shown in Figure 4A. Likewise, our regression analysis shows a larger cost effectiveness advantage for intramural awards in clinical citations, and a smaller but still significant advantage for extramural awards in number of papers and RCR (Supplemental Fig 4).

### Comparison of scores for human-focused research, animal research, and molecular/cellular research

Next, we assign scores to papers and projects based on the extent to which they can be classified as human research, animal research or molecular/cellular research. Again, we do this for projects of different durations: 1–3 years, 4–6 years, and 7–10 years. Both intramural and extramural projects have the highest average scores for human research (Figure 5 and Supplemental Figure 5). However, long-duration intramural projects tend to have lower human scores, while long-duration extramural projects tend to have higher human scores. For animal and molecular/cellular scores, the trend is reversed: long-duration intramural projects tend to have higher animal and molecular-cellular scores, while long-duration extramural projects tend to have lower scores. The possibility that longer-term intramural projects are more human-focused, which might explain the clinical citation comparative advantage with respect to the extramural program, is therefore inconsistent with the data.

## Discussion

Taken together, these results demonstrate comparative advantages for extramural and intramural funding mechanisms. In particular, extramural funding seems to excel at generating raw knowledge and facilitating its downstream flow. In contrast, intramural funding mechanisms seem to have a comparative advantage at generating research, basic or human-focused, that successfully informs downstream clinical research, aligning with the agency’s mission. This could potentially be attributed to the selection process for directly hiring scientists whose research agendas closely match the agency’s objectives, though this aspect wasn’t directly assessed in our study. However, we do rule out an obvious explanation: that more human-focused work in the intramural program is more likely to be conceptually closer to clinical trials and, therefore, have a lower barrier to entry into clinical studies (Kim, Levine, Nehl, & Walsh, 2020;

Weber, 2013). Intramural research is characterized by long-duration projects in contrast to the extramural portfolio, and these appear less human-focused than the extramural portfolio. Even when accounting for hidden costs borne by universities (Supplemental Figure 4), the fundamental pattern holds: intramural and extramural mechanisms have distinct comparative advantages that align with their institutional incentives. This suggests that these differences may be primarily driven by structural features, like mission alignment, of these funding mechanisms.

This study is not without limitations. First, data about the intramural portfolio is only available from post-2008, which constrains the time frame for this study. Second, collaboration between intramurally and extramurally funded scientists introduces complexity to the comparative analysis, leading to the exclusion of jointly funded publications. Third, the primary analysis uses an agency perspective based on NIH expenditures recorded in RePORTER, which does not capture the full economic cost of extramural research borne by universities. To address this, we conducted a sensitivity analysis (Supplemental Figure 4) inflating extramural indirect costs to account for unrecovered expenditures. Finally, recent changes at the NIH and the Department of Health and Human Services (which oversees the NIH), as well as changes proposed by the Senate, could lead to significant changes in the budget and organization of the NIH, and also in the processes it uses to make funding decisions.

Critiques of NIH’s grant review process often cite its conservatism, with a strong emphasis on preliminary data to mitigate project failure risks (Packalen & Bhattacharya, 2020). The recent creation of the Advanced Research Projects Agency for Health (ARPA-H) was in part for this reason (Collins, Schwetz, Tabak, & Lander, 2021). Although the high-risk, high-reward NIH portfolio seems to be largely effective at identifying and funding such projects (Tabak et al., 2019), its overall proportion of the total portfolio remains relatively small in favor of more traditional investigator-initiated research project grants. Because intramural researchers face retrospective rather than prospective review, this conservatism might be expected to manifest in a comparative advantage across a variety of measures for intramurally funded research. Competing theories suggest that extramural research may hold advantages on an investment-adjusted basis because of cost-sharing at universities, particularly for student labor (Zhang et al., 2022). Notably, intramural research focused on human/molecular or animal research seems to be particularly effective at generating clinically relevant research outputs (Supplemental Figure 5). Our findings reveal a nuanced reality: extramural institutions hold an edge in publication and citation rates aligned with their internal review procedures, while intramural research excels at stimulating bench-to-bedside translation on an investment-adjusted basis.

This analysis uses an agency-centered perspective to estimate comparative cost effectiveness, but there are other aspects of these two funding models that should be acknowledged (Drummond, Sculpher, Torrance, O’Brien, & Stoddart, 2005). First, while we estimated a more complete cost estimate through our modeling of a 30% indirect cost increase, this likely does not cover the complete degree of university investment in research. For example, startup costs are not modeled, nor are university investments that come from alternative funding streams like philanthropy, endowments, or state budgetary contributions. Extramural investigators also have hidden costs in the form of time spent preparing and managing grant applications, a cost that is not shared by intramural researchers. This represents an unreimbursed cost to the extramural funding system (Ioannidis, Gross, & Bergstrom, 2019), although this cost should be reflected in our productivity measures. Thus, our estimates of the comparative cost effectiveness of intramural research may be conservative. However, these alternative contributions to extramural research, which take the form of graduate student training (Zhang et al., 2022), faculty retention and infrastructure development (*Research Universities and the Future of America*, 2012), have real societal impacts. While the intramural program does train a limited number of graduate students, its postdoctoral workforce is its primary focus. Thus the extramural program’s contributions to a more highly educated United States workforce pipeline should be taken into consideration as a strength alongside its research outputs.

The distinction between the NIH intramural and extramural programs is sometimes controversial. A 2024 Senate proposal from the Senate Health, Education, Labor, and Pensions Committee suggested reforms to both the extramural and intramural programs (Cassidy, 2024). Feedback from a Request for Information suggested that respondents believe that these two programs are in many cases functionally equivalent. One salient consideration was how to best differentiate the extramural and intramural portfolios. Recommended changes to the extramural program focused on stimulating innovation and encouraging high-risk, high-reward projects. Proposals for the intramural program focused on increasing collaboration among funding institutes, promoting replication studies, and introducing temporary rotating intramural investigators focused on addressing long term, unmet needs. Both programs were recommended to increase focus on basic research. NIH was also encouraged to improve transparency by providing the public with more data to conduct metascience studies, in order to learn from research failures and build on scientific successes.

The present work suggests two avenues to build on the comparative strengths of intramural and extramural funding mechanisms. First, although the Senate proposal seems to indicate that the NIH as a whole refocus on basic research, it is the extramural program that seems comparatively well positioned for this task. Federal expenditures per article and downstream knowledge flow are lower in the extramural portfolio. By contrast, the intramural program is well-positioned to build on its strengths informing clinical research during structural reforms, if the most effective parts of the research portfolio are targeted. Supplemental Figure 5 suggests that animal research or research that combines aspects of human, animal, and molecular/cellular biology simultaneously within this portfolio are particularly cost effective at this goal. This aligns well with a focus from the Senate proposal on interdisciplinary research, and these kinds of projects can be readily identified with the data that are already available. However, longer-term intramural projects show a decreasing correlation with human research activities (Figure 5). This trend may need to be reversed if the strengths in the intramural program are to be effectively aligned with long-term research horizons.

We find that the NIH extramural program is comparatively more cost effective at generating research products that are aligned with traditional markers of academic achievement, publications and citations. The intramural program is comparatively more cost effective at generating research outputs that align with the agency’s stated mission, in this case, research that informs downstream clinical research. If this result generalizes to other agencies, it has implications for their portfolio management. The Department of Defense has a sizeable research portfolio in the biomedical space, most notably funded directly through the department itself or through the Congressionally Mandated Research Priorities program (“Congressionally Directed Medical Research Programs,” 2025). These results imply that refocusing that research portfolio with a focus on basic research in the extramural portfolio and health-informing research in the intramural portfolio may build on the programs’ respective incentive structures. However, the Department of Defense as well as other agencies like the National Science Foundation’s Federally Funded Research and Development Centers (FFRDCs) (“Federally Funded R&D Centers,” 2026) or the Department of Energy’s combination of national labs and FFRDCs may benefit from a refocus of their intramural programs toward research focused on advancing applied outcomes pertaining to their portfolios. If institutional incentive alignment is a significant contributor to the findings here, then these results are likely to be reflected in agencies whether or not their applied research goals are health-oriented.

## Methods

### Data

We collected the original NIH project data from NIH RePORTER (“NIH ExPORTER,” 2025), which contains 433,930 projects with funding information spanning from 1985 to 2019. We identified projects’ activity categories by looking up their first three letters in the project number (activity code). We classified the projects into intramural and extramural projects by the initial letter of their activity codes. Specifically, projects with an activity code starting with ‘Z’ were intramural projects, and other projects were extramural projects. Using this strategy, we identified 9,225 intramural projects and 424,705 extramural projects in the raw dataset. We retrieved the publication records for these projects by PMID indexed by PubMed (Medicine, 2020). These include mostly journal articles, although a small number of NIH-funded preprints are included as well (Funk, Zayas-Caban, & Beck, 2024; Hong, Hutchins, & Ni, 2026; Lindsay Nelson et al., 2022; L. Nelson et al., 2022).

The data cleaning process is as follows. Since the scientific focus of a study may drift over time, we dropped 70,297 projects with renewal records in our data. Second, considering intramural projects might change their activity categories and the three initial project number letters (e.g., ZIA changed to ZIH), we normalized 3,105 intramural project numbers by matching the rest of the project numbers to avoid inconsistency. Third, to focus on activity categories intended as research-oriented and exclude practice-oriented activity categories, at the activity level, we selected activity categories where at least 75% of projects had produced at least one paper. This step kept 106 activity codes (including six intramural activity codes). Although this process was designed to rule out intramural projects that had start dates prior to the beginning of data collection at 2009, it is possible that some intramural projects had earlier start dates than the records indicate. For example, if there was a gap year of funding in 2008, our ability to detect these might be more limited, or for example if the grant serial number changed. This is a technical limitation to our approach.

In this study, we focus on projects initially funded after 2008 and select the ten years from 2009 to 2019 as our analysis period. A total of 122,815 projects fell in this period. To remove the rest of potential non-research-oriented projects, at the individual project level, we selected 1% of projects with the highest cumulative ratio of funding and publication number and 1% of projects with the lowest. We then trained a random forest model to predict the projects most likely to be non-research-oriented based on their title and abstracts. Based on the predicted probability, we excluded 5% of intramural (84) and extramural (5,347) projects. The final analytical sample consists of 98,648 projects, including 97,054 extramural projects and 1,594 intramural projects, which produced 621,138 papers during our time window. Papers with both intramural and extramural funding were excluded. However, papers with multiple intramural funding links or multiple extramural funding links were included; most papers acknowledged only one project (Supplemental Figure 7).

Topic level ratios were calculated as a ratio of the proportions of total grants a topic represented in the intramural vs. extramural portfolios:

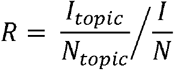

Where *I* is the number of intramural awards and *N* is the number of intramural plus extramural awards.

### Cost effectiveness

We used the primary project costs (direct costs, indirect costs for extramural institutions, and total costs, which combines these two) listed on NIH RePORTER as the cost data source. Subproject costs were not calculated. This approach takes an agency perspective, measuring cost effectiveness in terms of NIH expenditures. This does not capture the full investment in research expenditures at extramural institutions, so we also conduct a robustness check of our analyses where indirect costs are inflated by 30% (Supplemental Figure 4), estimated to be close to the unrecovered costs that universities invest (Droegemeier, 2017). Given the influence of price changes and inflations over years, we converted total costs (direct costs plus indirect costs) at the 2015 price level using NIH’s Biomedical Research and Development Price Index.

We used five paper-level metrics to measure the research output: number of papers; relative citation ratio (Arabi et al., 2025; B. I. Hutchins et al., 2017; B. I. Hutchins et al., 2016); approximate potential to translate (B. I. Hutchins, Davis, et al., 2019; Santangelo, 2017; Weber, 2013); total clinical citation counts (Travis A. Hoppe, Arabi, & Hutchins, 2023; B. Ian Hutchins, 2021; B. I. Hutchins, Baker, et al., 2019; “iCite,” 2015); and number of papers once received clinical citations. A project’s total research performance regarding a certain metric in one year is approximated as the sum of that metric for every paper published in that year.

Based on the project costs and research outputs, we calculated the cost per output as follows.

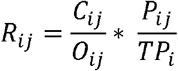

Where R_ij_ is the cost per output for project i in year j, C_ij_ the cumulative sum of funding costs for project i up to year j, O_ij_ is the cumulative sum of a certain research performance metric for project i up to year j, P_*ij*_ is the cumulative number of papers for project i up to year j, and *TP*_*i*_ is the total number of papers for project i until 2020.

### Regression analysis

We run the following regression model at the project level to estimate the differences between extra- and intramural projects for every year after the projects started.

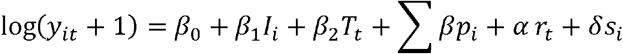

where *y*_*it*_ is project *i*’s deflated cost efficiency regarding a certain research performance *t* years after the funding start year; *I*_*i*_ is whether project *i* is an intramural project (1 if yes); *T*_*t*_ is the length of funding years until that time point, equal to the minimum of t and the total funding years; *p*_*i*_ stands for the PI-related variables; *r*_*t*_ and *s*_*i*_ are the year’s and project topic’s fixed effects. We transformed the *y*_*it*_ into log(*y*_*it*_ + 1) to mitigate the impact of uneven distribution.

PI-related variables include the number of past publications, number of past projects, number of PIs, share of clinical papers, past publications’ average relative citation ratio, publication experience, project experience, number of collaborators. We downloaded the PubMed Knowledge Graph datasets (Xu et al., 2020) to help extract the PI level variables. The dataset has disambiguated the authors of PubMed indexed publications and assigned unique identifiers to the authors. We matched both project numbers and paper author names with the datasets to find the PI’s assigned unique identifiers. We successfully retrieved the PI information for 98803 (94.9%) projects. The PI-related variables before the project funding started were extracted for every project, which played the proxy role of the input for the projects.

Another control variable, project topic, is calculated by performing a K-means clustering based on the NIH spending categories for all the projects in the sample. Project topic is defined by unsupervised clustering of semantic embeddings (Afshar, Yang, Thebault-Spieker, & Hutchins, 2026; Travis A. Hoppe, Arabi, & Hutchins, 2022; T. A. Hoppe et al., 2019) rather than graph clustering (Davis et al., 2025; Ni & Hutchins, 2025). To restrict the dimensionality, the 100 most frequent NIH spending categories were used in clustering, which cover 96.4% of all projects. We tried k=3, 4…10 and finally selected k=5 which generated the highest silhouette score.

As a robustness check, we used propensity score matching to reduce the potential confounding biases that may affect the outputs of interest and increase the comparability between intramural and extramural projects. For every year after the projects started, we used all project-level variables, including the length of funding years until that time point, PI-related variables, and project topics, to predict the propensity score of each project by fitting a logistic regression model. The propensity score shows the probability of a project to be an intramural project, based on the observable variables. For each intramural project, we selected one and four extramural projects, respectively, with the nearest propensity scores from a pool of extramural projects with the same funding start year. The regression model was run on the PSM sample again to check the robustness of previous results.

The PSM steps are as follows.

1. Estimate propensity scores. The study fits a propensity model (logistic) using covariates like productivity, collaboration, PI history, project duration, and project topic dummies. Multiple models are fit with balance checking for robust scoring.
2. Match within the same year: Treated and control records are paired only within the same year. For each treated project, the four closest controls are retained, and only very tight matches (diff < 0.001) are kept to enforce near-identical propensity scores.
3. Construct the matched cohort: Treated and control project IDs from the matched pairs are combined into a single dataset, forming the final matched sample.
4. Run outcome regressions on matched data: For each progression year, this study uses Stata reghdfe to run on log-transformed outcomes, with the project type as the key regressor and controls for productivity, collaboration, PI history, project duration, and project topic. The regressions contain fixed effects of project fiscal years.
5. Collect and visualize results: Coefficients, p-values, confidence bounds, and sample sizes are extracted per outcome and progression years, then plotted across progression years with shaded confidence bands and a zero reference line.

### Paper features

We used the concatenated documents of a paper’s title and abstract to train a word2vec model (Analysis, 2018; Analysis, Intelligence, & Institute, 2019) to classify the papers into clusters (Afshar et al., 2026; T. A. Hoppe et al., 2019). We removed common stop words, punctuation, and content lacking semantic information before training. During clustering, each paper’s document is represented as a 300-dimension vector by summing its each unique word’s vector weighted by its IDF. Principal component analysis (PCA) dimensionality reduction is applied to these 300-dimensional vectors to identify the 25 most influential components. We finally performed spectral clustering method using the document vectors and extract highly-frequent words to determine the cluster property and labels. Word2Vec nearest neighbor terms were uploaded to ChatGPT 3.5 to develop more human-readable labels.

## Supplemental Materials

### Supplemental Text

Questions about the most effective approaches to structure portfolio management for science funders have been a source of contention. This is primarily due to the conflicting priorities among government officials, the mission of funding agencies, and the perspectives of scientific researchers (Goldstein & Kearney, 2020). While the 2018 Evidence Act (Abraham & Haskins, 2017; Young, 2021) mandates that all science funders incorporate data-driven decision-making, the U.S. Congress played a significant role in catalyzing such efforts, particularly at the National Institutes of Health (NIH), through the establishment of divisions such as the Office of Portfolio Analysis in 2011 (Department, 2011). The division was created to advance these data-driven initiatives even before they were broadly implemented across other federal agencies. Consequently, NIH serves as an excellent case study for policy examination, given its more extensive and robust data infrastructure compared to other agencies.

A pressing question that often surfaces, particularly when facing inflation-adjusted budgetary declines, concerns the comparative efficiency of externally funded grants, usually awarded to universities, medical institutions, and research centers, in contrast to intramurally funded projects where scientists are employed directly as government personnel and conduct research within federal facilities. The 2013 sequester (Fox, 2013), a budget reduction mechanism that abruptly removed a sizeable fraction of government funding for scientific research, revealed significant contention in the scientific community about the extent to which extramural versus intramural funding should shoulder the burden of budgetary declines (Scientopia, 2014).

Various theories exist to highlight the respective merits of these two funding models. Extramural institutions are thought to engage in extensive cost-sharing that might reduce the degree of government investment necessary to stimulate scientific advancement (Culliton, 1992; Korn, 2015; Macilwain, 1999). In effect, despite the negotiated indirect costs that are paid to offset institutional overhead that supports scientific research at extramural institutions, these institutions often contribute additional resources that foster science advance. Research indicates that institutional contributions, particularly in terms of trainee labor (Zhang, Wapman, Larremore, & Clauset, 2022), are important for stimulating scientific productivity, supporting this theory. Moreover, it is essential to recognize the substantial portion of extramural funding typically dedicated to training students and early career researchers. This investment not only aids in producing the next generation of researchers in the field but also contributes to the long-term sustainability of the research workforce (Harris, 2014). On the other hand, the direct hiring of scientists by the government under the intramural funding model allows for the selection of researchers whose research agendas more closely align with the agency’s mission. Furthermore, despite shouldering the entire cost of intramural research, intramural PIs are freed from the time and resource burdens associated with grant applications. This freedom allows them to focus entirely on advancing scientific knowledge in their respective fields. Nonetheless, intramural grants may encounter constraints on autonomy due to their affiliation with a larger government institution, potentially restricting their freedom through manuscript clearance processes. This affiliation also implies that intramural researchers may not be entirely shielded from potential bureaucratic hurdles and unwarranted administrative burdens that can impede the progression of scientific endeavors. Therefore, each funding approach has unique strengths and considerations, making it essential to carefully weigh the advantages and disadvantages of extramural and intramural funding when allocating resources for scientific research.

### Supplemental Figures

**Supplemental Figure 1.**
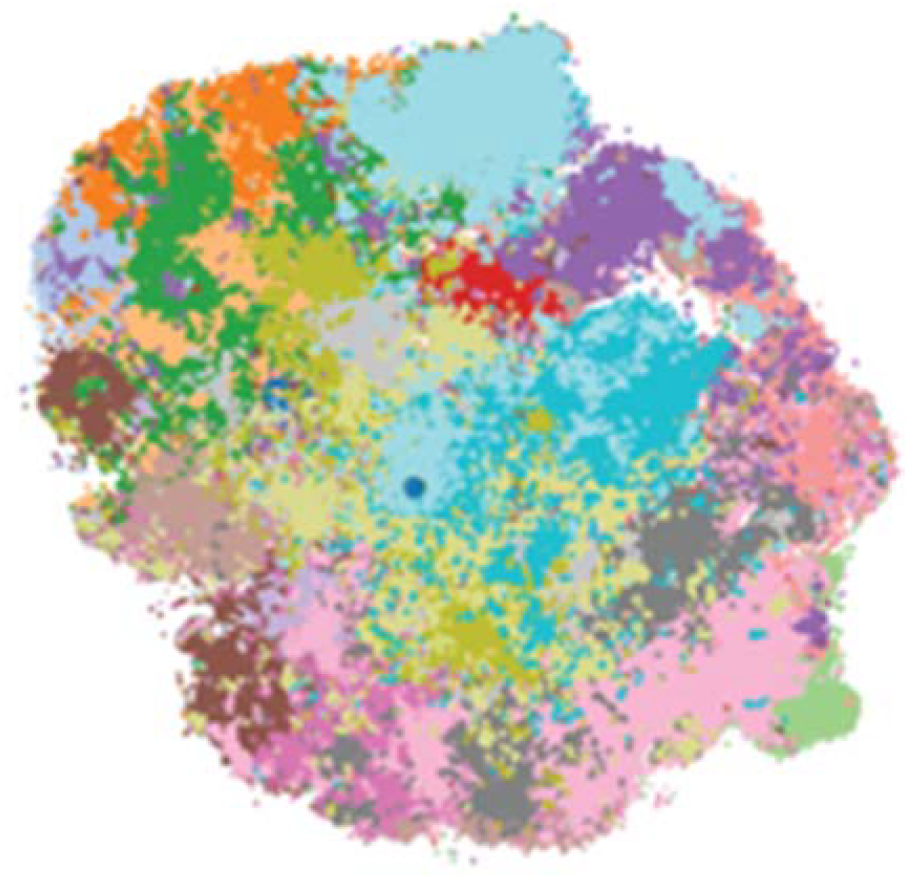
T-SNE plot illustrating the distribution of topic clusters, with colors consistent with those in Figure 1. This shows a dimensionality reduction of the locations of these grants in Word2Vec space, indicating their relative proximity to one another. Grants with similar semantic meaning appear closer to one another, while those with less semantic similarity will appear farther away. Similar grants tend to cluster together in T-SNE visualizations.

**Supplemental Figure 2.**
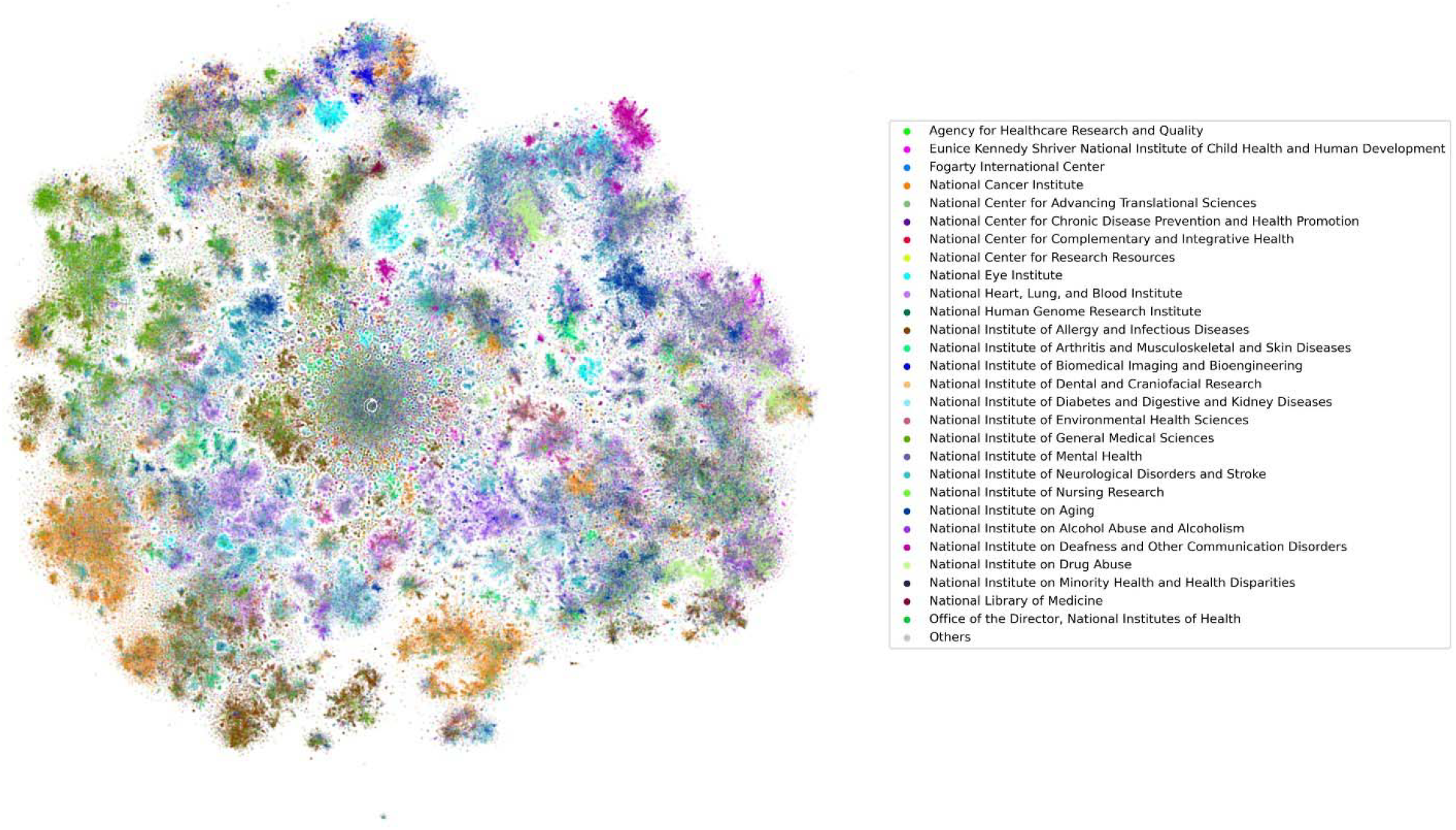
T-SNE plot illustrating the distribution of grants in NIH institutes, using the same methodology as Supplemental Figure 1. This shows a dimensionality reduction of the locations of these grants in Word2Vec space, indicating their relative proximity to one another. Grants with similar semantic meaning appear closer to one another, while those with less semantic similarity will appear farther away. Similar grants tend to cluster together in T-SNE visualizations.

**Supplemental Figure 3.**
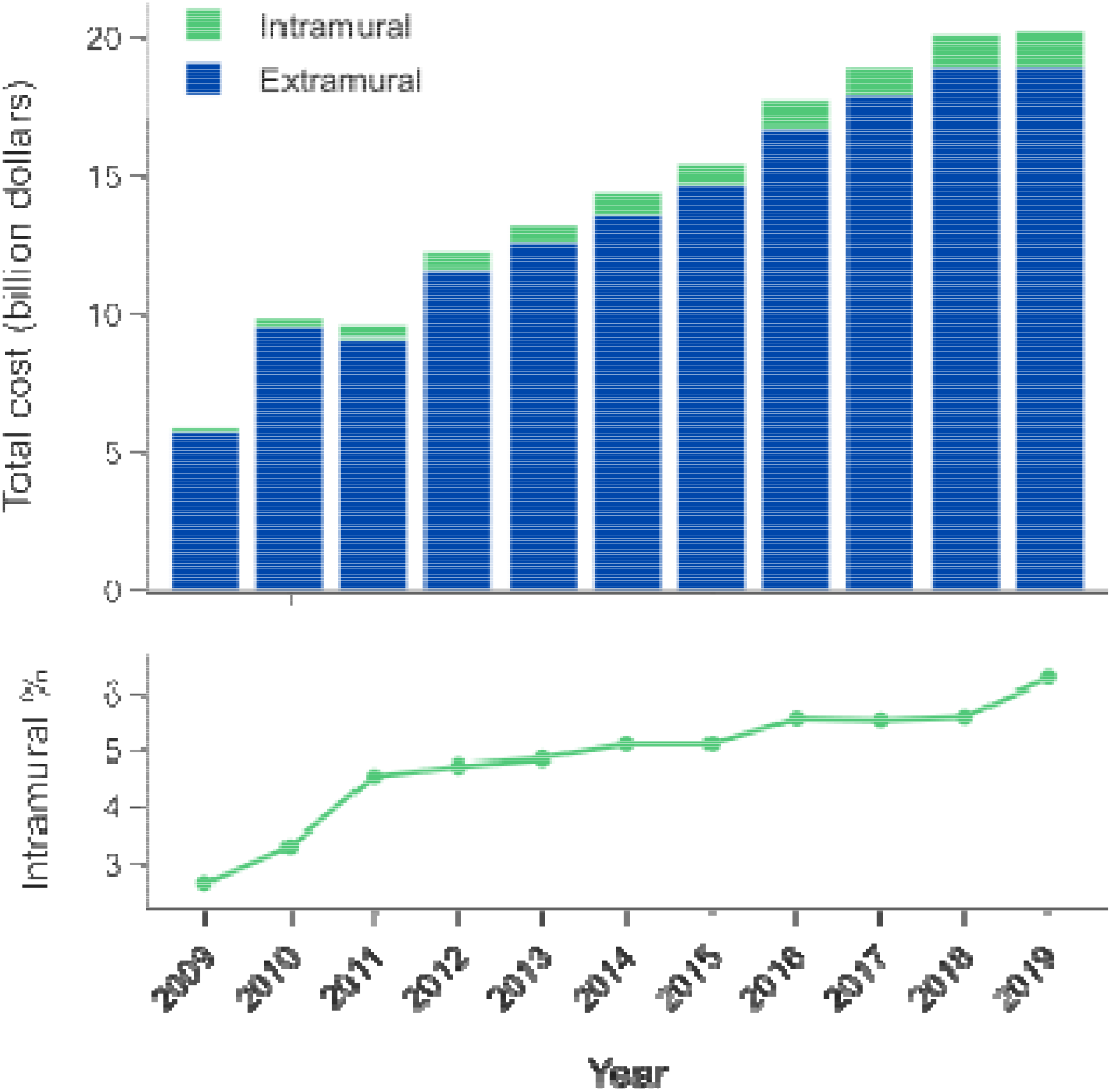
Cost breakdown for our sample of projects (see Methods) by Extramural or Intramural origin.

**Supplemental Figure 4.**
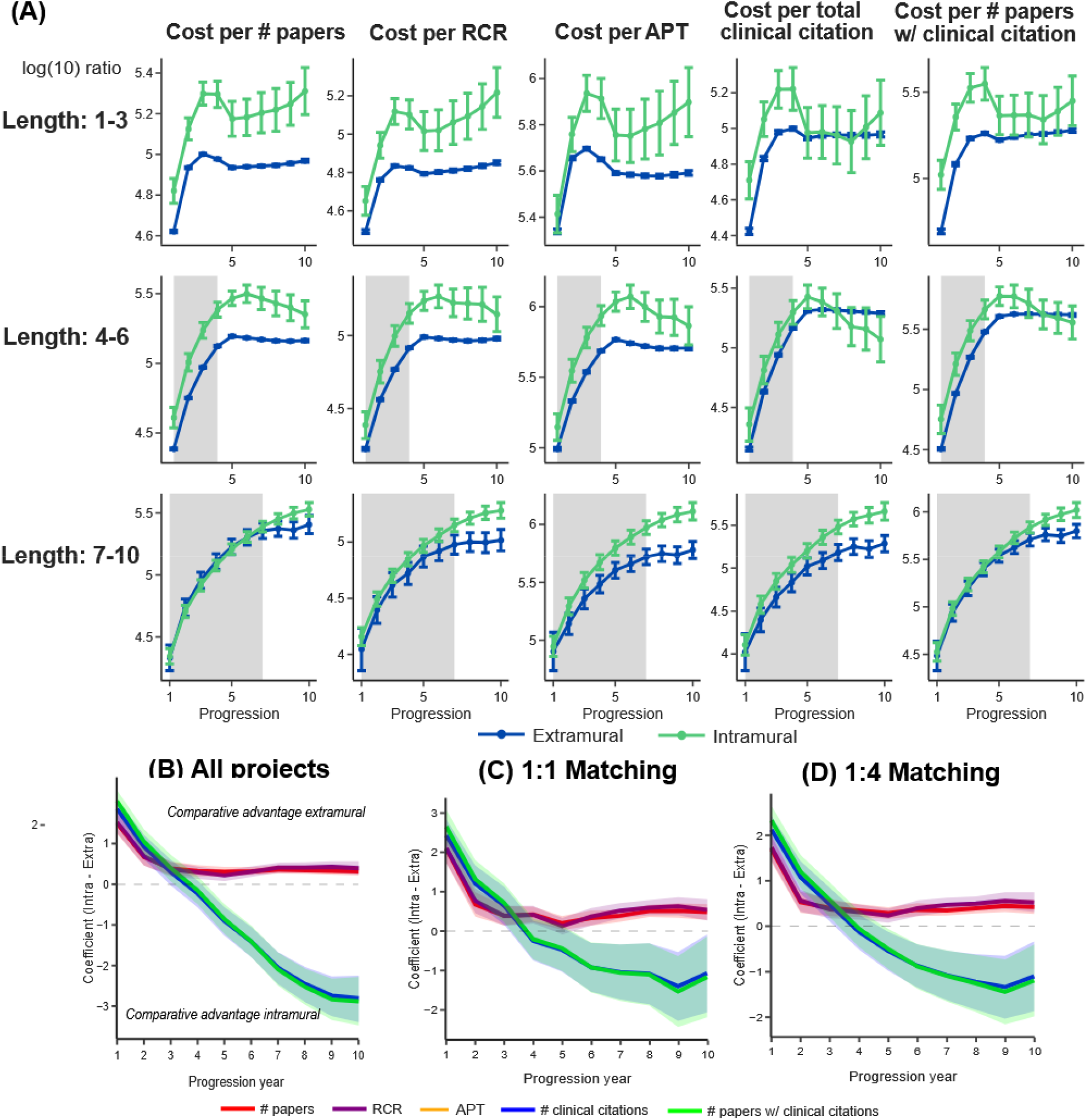
Cost effectiveness of intramural and extramural projects, with Indirect costs for Extramural projects inflated by 30%. (A) A measure of cost effectiveness versus progression (i.e., year of grant) for intramural research (green) and extramural research (red), for projects of different durations: 1–3 years (top row), 4–6 years (middle), and 7–10 years (bottom). These regressions do not control for other characteristics, but rather represent the raw ratios. For the first column, the Y-axis displays log10(ratio) +1, where ratio is the cumulative total costs to the cumulative total research output for each metric (cost:output, for the first column output = #papers); error bars denote the 95% confidence intervals. The remaining columns show measures of cost effectiveness for relative citation ratio, approximate potential to translate, total clinical citation counts, and a binary measure of clinical citations. To account for the fact that many papers are published after funding for the relevant grant has ended, grant amounts were multiplied by a deflator – this represents the proportion of papers published to date against the anticipated number of future publications, as determined by empirical measurements (Supplemental Table 1). In most cases, according to this analysis, extramural research is more cost effective than intramural research when observing uncontrolled regressions. (B–D) Linear regression results of the cost efficiency of research output measures against project types (intramural vs. extramural). The regression model was fitted for each year of the project’s progression. Unlike panel (A), this regression model controls for grant, investigator, and collaboration characteristics in order to obtain a more accurate estimate of the relative cost efficiency of intramural vs. extramural projects. The Y-axis coefficient indicates the mean disparity in research output between intramural and extramural projects, controlling for these other variables (see Methods). Because there might be covariates that could confound the data, separate regressions were conducted for all projects (B, the default), and for balanced projects using 1:1 propensity score matching (1 extramural grant for every 1 intramural grant) in order to compare grants that were the most similar to reduce the influence of unobserved covariates (C) and (D) similarly to (C) 1:4 propensity matching as a robustness check.

**Supplemental Figure 5.**
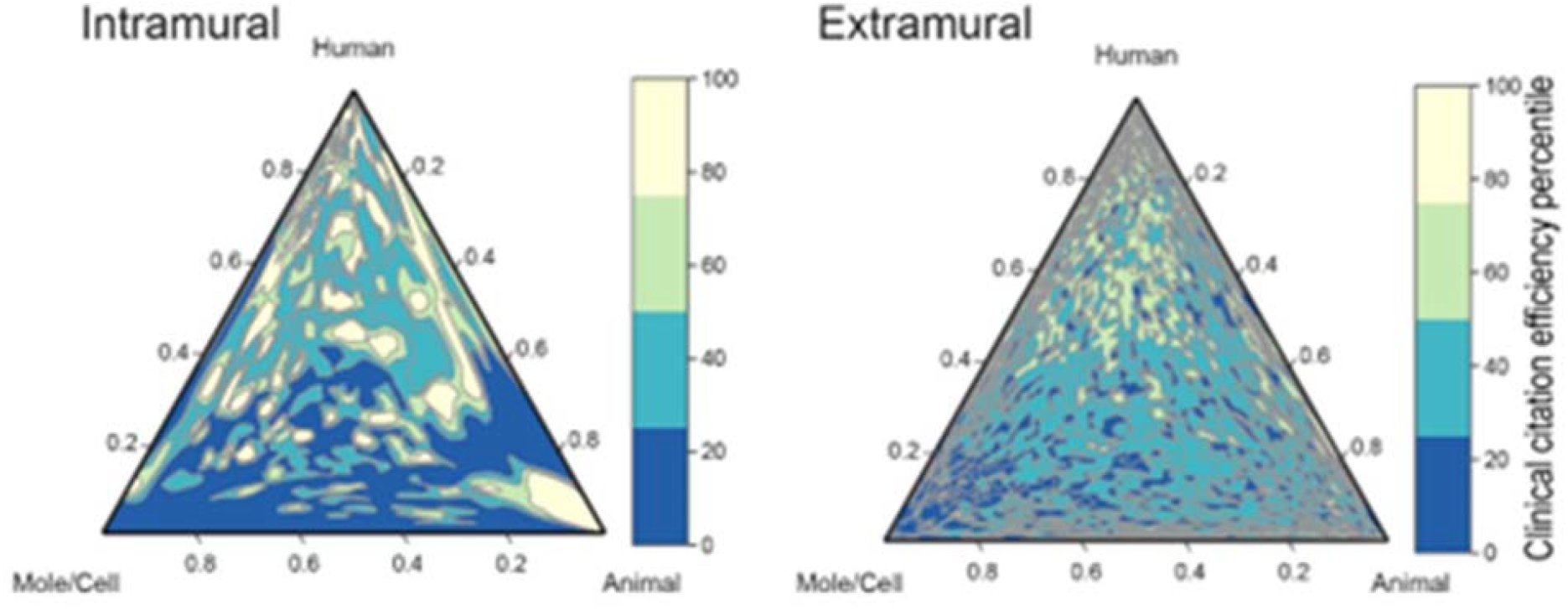
Ternary contour plots representing the clinical citation efficiency in the human, animal, and molecular/cellular score system for intramural (D) and extramural projects (E). (Hutchins, Davis, Meseroll, & Santangelo, 2019; Weber, 2013). Here, efficiency was the percentile of the cost per output in descending order. Each contour line denotes a constant efficiency percentile. Yellow/green are high-efficiency areas of the triangle, and blues are low-efficiency areas.

**Supplemental Figure 6.**
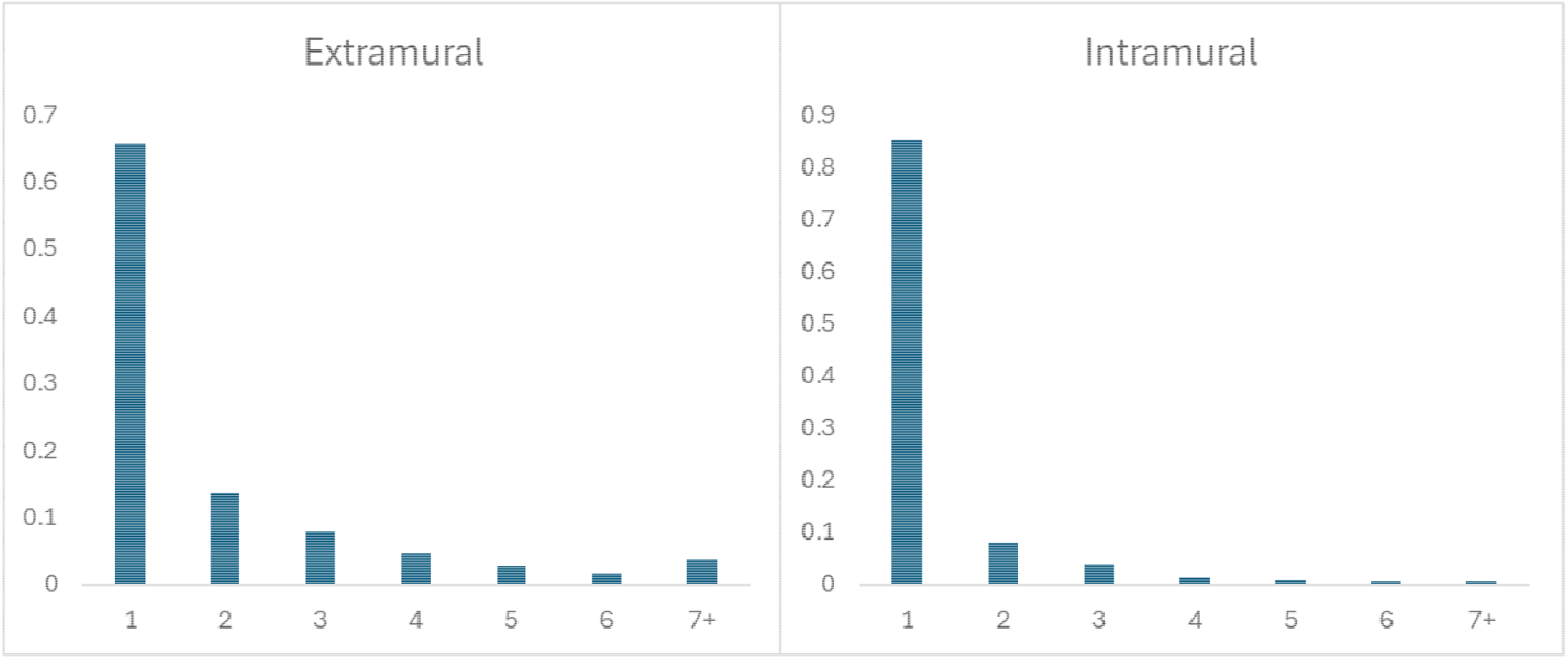
Distribution of papers acknowledging more than one funding source, divided, Extramural and Intramural.

**Supplemental Table 1.** Number and proportion of papers published after grant ended by activity code prefix.

| Activity code prefix | # Papers after funding ended | # Total papers | % Funded papers published after grant ended |
| --- | --- | --- | --- |
| D | 9,213 | 13,039 | 70.66% |
| F | 18,649 | 32,097 | 58.10% |
| G | 283 | 1,095 | 25.84% |
| K | 43,633 | 102,892 | 42.41% |
| P | 20,766 | 50,650 | 41.00% |
| R | 225,490 | 418,989 | 53.82% |
| S | 1,061 | 2,176 | 48.76% |
| T | 5,324 | 12,839 | 41.47% |
| U | 49,222 | 112,459 | 43.77% |
| Z | 1,794 | 19,871 | 9.03% |

## References

Afshar, A. S., Yang, Q., Thebault-Spieker, J., & Hutchins, B. I. (2026). Quantifying (mis)alignment between reader focus and editor citation in scholarly biomedical topics in Wikipedia. Quantitative Science Studies, 1–23. doi:10.1162/QSS.a.457

Analysis, N. I. o. H. O. o. P. (2018). National Institutes of Health word2vec pipeline. Retrieved from https://github.com/NIHOPA/word2vec_pipeline

Analysis, N. I. o. H. O. o. P., Intelligence, L., & Institute, T. A. R. (2019). Natural Language Preprocessing (NLPre) 2.0. GitHub. Retrieved from https://github.com/NIHOPA/NLPre

Arabi, S., Ni, C., & Hutchins, B. I. (2025). Most researchers would receive more recognition if assessed by article-level metrics than by journal-level metrics. PLoS Biol, 23(12), e3003532. doi:10.1371/journal.pbio.3003532

Azoulay, P., Zivin, J. S. G., & Manso, G. (2009). Incentives and creativity: evidence from the academic life sciences. RAND J. Econ, 42, 527–554. doi:10.3386/w15466

Cassidy, B. (2024). NIH in the 21st Century: Ensuring Transparency and American Biomedical Leadership. Retrieved from https://www.help.senate.gov/imo/media/doc/nih_modernization_5924pdf.pdf

Collins, F. S., Schwetz, T. A., Tabak, L. A., & Lander, E. S. (2021). ARPA-H: Accelerating biomedical breakthroughs. Science, 373(6551), 165–167. doi:10.1126/science.abj8547

Congressionally Directed Medical Research Programs. (2025). Retrieved from https://cdmrp.health.mil/

Culliton, J. J. (1992). “Indirect” Costs Are Real Costs at Massachusetts Institute of Technology. Science, 255(5046), 778–779. doi:10.1126/science.1535998

Davis, M. T., Busse, B. L., Arabi, S., Meyer, P., Hoppe, T. A., Meseroll, R. A., Santangelo, G. M. (2025). Prediction of transformative breakthroughs in biomedical research. bioRxiv. doi:10.64898/2025.12.16.694385

Droegemeier, K. (2017). Testimony before the House Appropriations Subcommittee on Labor, Health and Human Services, Education, and Related Agencies. Retrieved from https://docs.house.gov/meetings/AP/AP07/20171024/106525/HHRG-115-AP07-Wstate-DroegemeierK-20171024.pdf

Drummond, M. E., Sculpher, M. J., Torrance, G. W., O’Brien, B. J., & Stoddart, G. L. (2005). Methods for the Economic Evaluation of Health Care Programmes.

Federally Funded R&D Centers. (2026). Retrieved from https://ncses.nsf.gov/resource/master-gov-lists-ffrdc

Funk, K., Zayas-Caban, T., & Beck, J. (2024). Phase 1 of the National Institutes of Health Preprint Pilot: Testing the viability of making preprints discoverable in PubMed Central and PubMed. bioRxiv. doi:10.1101/2022.12.12.520156

Goldstein, A. P., & Kearney, M. (2020). Know when to fold ‘em: An empirical description of risk management in public research funding. Research Policy, 49(1). doi:ARTN 103873 10.1016/j.respol.2019.103873

Hong, X., Hutchins, B. I., & Ni, C. (2026). Faster science, penalties in evaluation, and concerns on quality and impact: Researchers’ use and perceptions of preprints. bioRxiv. doi:10.64898/2026.03.02.709147

Hoppe, T. A., Arabi, S., & Hutchins, B. I. (2022). Predicting causal citations without full text. bioRxiv. doi:10.1101/2022.07.05.498860

Hoppe, T. A., Arabi, S., & Hutchins, B. I. (2023). Predicting substantive biomedical citations without full text. Proceedings of the National Academy of Sciences, 120(30). doi:10.1073/pnas.2213697120

Hoppe, T. A., Litovitz, A., Willis, K. A., Meseroll, R. A., Perkins, M. J., Hutchins, B. I., Santangelo, G. M. (2019). Topic choice contributes to the lower rate of NIH awards to African-American/black scientists. Sci Adv, 5(10), eaaw7238. doi:10.1126/sciadv.aaw7238

Hutchins, B. I. (2021). A tipping point for open citation data. Quantitative Science Studies, 1–5. doi:10.1162/qss_c_00138

Hutchins, B. I., Baker, K. L., Davis, M. T., Diwersy, M. A., Haque, E., Harriman, R. M., Santangelo, G. M. (2019). The NIH Open Citation Collection: A public access, broad coverage resource. PLoS Biol, 17(10), e3000385. doi:10.1371/journal.pbio.3000385

Hutchins, B. I., Davis, M. T., Meseroll, R. A., & Santangelo, G. M. (2019). Predicting translational progress in biomedical research. PLoS Biol, 17(10), e3000416. doi:10.1371/journal.pbio.3000416

Hutchins, B. I., Hoppe, T. A., Meseroll, R. A., Anderson, J. M., & Santangelo, G. M. (2017). Additional support for RCR: A validated article-level measure of scientific influence. PLoS Biol, 15(10), e2003552. doi:10.1371/journal.pbio.2003552

Hutchins, B. I., Yuan, X., Anderson, J. M., & Santangelo, G. M. (2016). Relative Citation Ratio (RCR): A New Metric That Uses Citation Rates to Measure Influence at the Article Level. PLoS Biol, 14(9), e1002541. doi:10.1371/journal.pbio.1002541

iCite. (2015). Retrieved from https://icite.od.nih.gov/

iCite, Hutchins, B. I., & Santangelo, G. M. (2019). iCite Database Snapshots (NIH Open Citation Collection). Retrieved from: 10.35092/yhjc.c.4586573

Ioannidis, J. P. A., Gross, K., & Bergstrom, C. T. (2019). Contest models highlight inherent inefficiencies of scientific funding competitions. PLOS Biology, 17(1). doi:10.1371/journal.pbio.3000065

Kim, Y. H., Levine, A. D., Nehl, E. J., & Walsh, J. P. (2020). A Bibliometric Measure of Translational Science. Scientometrics, 125(3), 2349–2382. doi:10.1007/s11192-020-03668-2

Korn, D. (2015). Indirect costs: Cash is no gravy train. Nature, 517(7535), 438–438. doi:10.1038/517438b

Lauer, M., Roychowdhury, D., Patel, K., Walsh, R., & Pearson, K. (2017). Marginal Returns and Levels of Research Grant Support among Scientists Supported by the National Institutes of Health. bioRxiv. doi:10.1101/142554

Macilwain, C. (1999). NSF told to ease up cost-share demands. Nature, 399(6732), 95–95. doi:10.1038/20039

Medicine, N. L. o. (2020). Download MEDLINE/PubMed Data. Retrieved from https://www.nlm.nih.gov/databases/download/pubmed_medline.html

National Cancer Institute (NCI) Center for Cancer Research. (2025). Retrieved from https://www.nih.gov/about-nih/nih-almanac/national-cancer-institute-nci

Nelson, L., Ye, H., Schwenn, A., Lee, C., Arabi, S., & Hutchins, B. I. (2022). Robustness of evidence reported in preprints during peer review. Research Square. doi:10.21203/rs.3.rs-1344293/v1

Nelson, L., Ye, H., Schwenn, A., Lee, S., Arabi, S., & Hutchins, B. I. (2022). Robustness of evidence reported in preprints during peer review. Lancet Glob Health, 10(11), e1684–e1687. doi:10.1016/S2214-109X(22)00368-0

Ni, C., & Hutchins, B. I. (2025). Framework for assessing the risk to a field from fraudulent researchers: A case study of Alzheimer’s disease. Journal of the Association for Information Science and Technology, 76(9), 1162–1173. doi:10.1002/asi.25009

NIH Budget. (2025). Retrieved from https://www.nih.gov/about-nih/organization/budget

NIH ExPORTER. (2025). Retrieved from https://reporter.nih.gov/exporter

Packalen, M., & Bhattacharya, J. (2020). NIH funding and the pursuit of edge science. Proceedings of the National Academy of Sciences, 117(22), 12011–12016. doi:10.1073/pnas.1910160117

Research Universities and the Future of America. (2012).

Sampat, B. N. (2012). Mission-oriented biomedical research at the NIH. Research Policy, 41(10), 1729–1741. doi:10.1016/j.respol.2012.05.013

Santangelo, G. M. (2017). Article-level assessment of influence and translation in biomedical research. Mol Biol Cell, 28(11), 1401–1408. doi:10.1091/mbc.E16-01-0037

Tabak, L., Lee, B., Carpten, J., Carnes, M., Miller, B., Palin, A., Valantine, H. (2019). Report of the ACD Working Group on High-Risk, High-Reward Research. Retrieved from https://acd.od.nih.gov/documents/presentations/06132019HRHR_B.pdf

Wahls, W. P. (2018a). Diminishing marginal returns on NIH grant funding to institutions. bioRxiv. doi:10.1101/367847

Wahls, W. P. (2018b). The NIH must reduce disparities in funding to maximize its return on investments from taxpayers. eLife, 7. doi:10.7554/eLife.34965

Weber, G. M. (2013). Identifying translational science within the triangle of biomedicine. J Transl Med, 11, 126. doi:10.1186/1479-5876-11-126

What happens to your application during and after review? (2025). Retrieved from https://public.csr.nih.gov/ForApplicants/InitialReviewResultsAndAppeals/applicationduringafterreview

Xu, J., Kim, S., Song, M., Jeong, M., Kim, D., Kang, J., Ding, Y. (2020). Building a PubMed knowledge graph. Scientific Data, 7(1). doi:10.1038/s41597-020-0543-2

Zhang, S., Wapman, K. H., Larremore, D. B., & Clauset, A. (2022). Labor advantages drive the greater productivity of faculty at elite universities. Science Advances, 8(46). doi:10.1126/sciadv.abq7056

## Supplemental References

Abraham, K. G., & Haskins, R. (2017). The Promise of Evidence-Based Policymaking. Retrieved from https://bipartisanpolicy.org/download/?file=/wp-content/uploads/2019/03/Full-Report-The-Promise-of-Evidence-Based-Policymaking-Report-of-the-Comission-on-Evidence-based-Policymaking.pdf

Department, H. a. H. S. (2011). National Institutes of Health Statement of Organization, Functions, and Delegations of Authority. Federal Register: Library of Congress Retrieved from https://www.federalregister.gov/documents/2011/02/09/2011-2848/national-institutes-of-health-statement-of-organization-functions-and-delegations-of-authority

Fox, J. L. (2013). Sequestration to slash research grants and delay biosimilars. Nature Biotechnology, 31(4), 271–271. doi:10.1038/nbt0413-271

Harris, A. (2014). Young, Brilliant, and Underfunded. New York Times. Retrieved from https://www.nytimes.com/2014/10/03/opinion/young-brilliant-and-underfunded.html

Scientopia. (2014). Berg posts data on NIH Intramural funding. Retrieved from https://drugmonkey.scientopia.org/2014/02/05/berg-posts-data-on-nih-intramural-funding/

Young, S. D. (2021). Evidence-Based Policymaking: Learning Agendas and Annual Evaluation Plans. (M-21-27). Retrieved from https://www.whitehouse.gov/wp-content/uploads/2021/06/M-21-27.pdf

